# Pharmacological activation of PIEZO1 in human red blood cells prevents *Plasmodium falciparum* invasion

**DOI:** 10.1101/2021.05.28.446171

**Authors:** Rakhee Lohia, Jordy Le Guet, Laurence Berry, Hélène Guizouarn, Roberto Bernal, Rachel Cerdan, Manouk Abkarian, Dominique Douguet, Eric Honoré, Kai Wengelnik

## Abstract

An inherited gain-of–function variant (E756 del) in the mechanosensitive cationic channel *PIEZO1* was recently shown to confer a significant protection against severe malaria. Here, we demonstrate *in vitro* that human red blood cell (RBC) infection by *Plasmodium falciparum* is prevented by the pharmacological activation of PIEZO1. The PIEZO1 activator Yoda1 inhibits RBC invasion, without affecting parasite intraerythrocytic growth, division or egress. RBC dehydration, echinocytosis and intracellular Na^+^/K^+^ imbalance are unrelated to the mechanism of protection. Inhibition of invasion is maintained, even after a prolonged wash out of Yoda1. Similarly, the chemically unrelated activators Jedi1 and Jedi2 potently inhibit parasitemia, further indicating a PIEZO1-dependent mechanism. Notably, Yoda1 treatment significantly reduced RBC surface receptors of *P. falciparum*, and decreased merozoite attachment and subsequent RBC deformation. Altogether these data indicate that the pharmacological activation of Piezo1 in human RBCs inhibits malaria infection by impairing *P. falciparum* invasion.

## Introduction

Malaria is caused by the cyclic infection and destruction of red blood cells (RBCs) by a protozoan parasite of the genus *Plasmodium*, the most deadly form being caused by *P. falciparum*. There is substantial variability in susceptibility to malaria in the human population and in numerous cases this has been attributed to particular features of human RBCs. Sickle cell anaemia, caused by a point mutation in haemoglobin (Hb) leading to the production of HbS, is the most well-known condition conferring resistance to malaria (****Allison, 1954****). Although having a drastic impact on well-being and survival, there is still continuous selection for these mutations in malaria endemic countries (***Elguero et al., 2015***). In addition, many other variations in RBC proteins and physiology were suggested to be linked to reduced susceptibility to malaria, including RBCs dehydration caused by a variety of clinical or experimental conditions (***Roberts and Williams, 2003; Tiffert et al., 2005***).

PIEZO1 and PIEZO2 are mechanosensitive non-selective cationic channels, permeable to Na^+^, K^+^ and Ca^2+^ (****Coste et al., 2010****). PIEZO1 is activated by a variety of mechanical stress, including local membrane stretch, whole cell indentation or shear stress (****Murthy et al., 2017****). PIEZO1 and PIEZO2 isoforms, which share 43% sequence identity, are conserved during evolution from plants, worm, fly, fish to human (****Coste et al., 2010****), while no orthologue is present in *Plasmodium* species or other apicomplexan parasites (****Prole and Taylor, 2013****). PIEZO1 and PIEZO2 are differentially expressed across different tissues and cell types, with only PIEZO1 present in RBCs (***Cahalan et al., 2015; Coste et al., 2010***). PIEZO1 and PIEZO2 are made of a remarkably large homotrimeric complex (about 900 kDa, including 114 transmembrane segments) with a curved shape, similar to a three blades propeller (or a triskelion)(***Ge et al., 2015; Guo and MacKinnon, 2017; Saotome et al., 2018; Wang et al., 2019; Zhao et al., 2018***). Recent findings indicate that PIEZO1 curves the membrane, forming an inverted cone that reversibly flattens upon activation in response to mechanical stimulation (***Guo and MacKinnon, 2017; Haselwandter and MacKinnon, 2018; Lin et al., 2019***).

The published proteome of the human RBCs indicates the presence of about 1000-1500 PIEZO1 channels per RBC (****Bryk and Wisniewski, 2017****). Gain-of-function (GOF) *PIEZO1* mutations causing a delayed inactivation (i.e. prolonged opening), are linked to hereditary xerocytosis, a mild haemolytic anaemia associated with RBCs dehydration (***Albuisson et al., 2013; Andolfo et al., 2013; Bae et al., 2013; Zarychanski et al., 2012***). Remarkably, mice carrying a xerocytosis mutation (mR2482H) either in all hematopoietic cells or specifically in erythrocytes show RBCs dehydration and are protected from cerebral malaria caused by the rodent-specific parasite *Plasmodium berghei* (***Ma et al., 2018; Nguetse et al., 2020***). *PIEZO1* R2482H mice show no damage of the blood-brain-barrier (BBB), while wild type mice succumb to cerebral malaria leading to death within 5 to 7 days post infection. Along this line, another GOF *PIEZO1* variant (E756 del), present in about 1/3 of the black population from Western Africa, also confers a significant resistance against severe malaria (***Ma et al., 2018; Nguetse et al., 2020***). Recently, it was confirmed that in children from Gabon, the *PIEZO1* E756del variant is strongly associated with protection against severe malaria in heterozygotes, independent of the protection conferred by the sickle cell trait (HbS) (****Nguetse et al., 2020****). Whether or not the protective effect of *PIEZO1* E756 del is linked to RBCs dehydration and a decreased intracellular growth of *P. falciparum* is disputed (***Ma et al., 2018; Nguetse et al., 2020***). Instead, surface expression of the *P. falciparum* virulence protein PfEMP-1, that mediates cytoadherence of infected RBCs to the endothelium in cerebral malaria, was shown to be significantly reduced in infected cells heterozygous for *PIEZO1* E756 (****Nguetse et al., 2020****). Moreover, additional evidence indicates that the beneficial effect of *PIEZO1* R2482H is not solely due to an effect on RBCs, but also involves the immune system (****Ma et al., 2018****). *PIEZO1* E756 has also been shown to lead to iron overload by increasing RBC turnover through phagocytosis by macrophages; however this effect is unrelated to malaria protection (***Ma et al., 2018; Ma et al., 2021***). All together these data indicate that PIEZO1 GOF is protective against severe malaria, although the mechanisms implicated still remain unclear (***Ma et al., 2018; Nguetse et al., 2020***).

The pharmacology of PIEZO1 is still at its early ages. Yoda1 was identified by a high throughput screening of small molecules as an activator of mouse and human PIEZO1, without affecting PIEZO2 (****Syeda et al., 2015****). When analysed in HEK 293T cells expressing either mouse or human PIEZO1, Yoda1 (in the micromolar range) causes enhanced opening of PIEZO1 in response to mechanical stimulation, together with a delayed inactivation (****Syeda et al., 2015****). Yoda1 acts as an activator of PIEZO1 (not as a true opener) by shifting its pressure-effect curve towards lower pressure values (****Syeda et al., 2015****). Two additional compounds Jedi1 and Jedi2 (chemically unrelated to Yoda1) similarly activate PIEZO1, although at a concentration in the millimolar range (****Wang et al., 2018****).

In mouse RBCs, Yoda1 causes a robust increase in intracellular calcium, without the requirement of a mechanical stimulation (****Cahalan et al., 2015****). Notably, addition of 15 μM Yoda1 to RBCs leads to a dramatic change in cell shape, inducing echinocytosis within 30 seconds (****Cahalan et al., 2015****). This change in cell shape is strictly dependent on the presence of PIEZO1 since RBC from knock out (KO) mice are not altered by Yoda1 (****Cahalan et al., 2015****). Yoda1 induces mouse RBC dehydration through the secondary activation of the Ca^2+^-dependent Gardos (KCNN4 / KCa3.1) channel and consequent water loss by aquaporins (***Cahalan et al., 2015; Rapetti-Mauss et al., 2017***).

Although the link between inherited *PIEZO1* GOF variants and resistance against malaria is now well established, the mechanism(s) of protection remain elusive and controversial (***Ma et al., 2018; Nguetse et al., 2020***). In this study, taking advantage of an *in vitro* model of human RBC infection by *P. falciparum*, we aimed at exploring the antimalarial activity of PIEZO1 pharmacological activators and at determining which step of the infection cycle is PIEZO1 sensitive. Altogether, our findings show that the *in vitro* pharmacological activation of PIEZO1 in human RBCs prevents parasite invasion, independently of RBC dehydration, Na^+^/K^+^ imbalance or echinocytosis. PIEZO1 activation reduced RBC surface receptors of *P. falciparum*, merozoite attachment and subsequent RBC deformation associated with the inhibition of parasite invasion.

## Results

### The PIEZO1 activator Yoda1 inhibits *P. falciparum* parasitemia *in vitro*

To evaluate *in vitro* the effect of PIEZO1 activation on *P. falciparum* growth in human RBCs, synchronized ring stage parasites were incubated for 60 h in the presence of different concentrations of Yoda1. Notably, a dose-dependent decrease in parasitemia was observed with an IC50 value of about 500 nM (***Figure 1A***).

**Figure 1.**
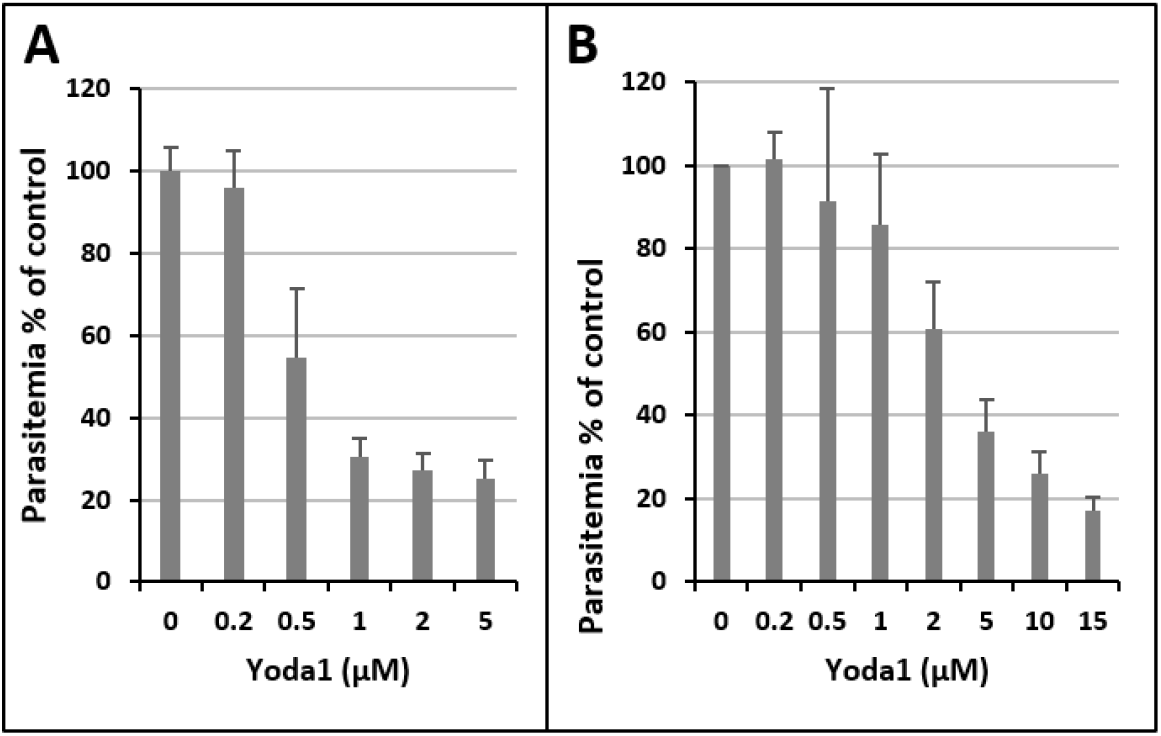
Pharmacological activation of PIEZO1 reduces *P. falciparum* parasitemia *in vitro*. (**A**) Synchronized ring stage parasite cultures in complete medium were incubated for 60 h in the continuous presence of Yoda1 at different concentrations before determining the parasitemia by FACS analysis (n=5). (**B**) Uninfected RBCs were first pre-treated in RPMI with different concentrations of Yoda1 for 10 min and then washed extensively before adding purified schizont/segmenter stage infected RBCs for culture in complete medium. Parasitemia was determined after incubation for 18 h (n= 3). Shown are mean values + SD. Kruskal-Wallis test for (A) p<0.001, (B) p<0.01.

Although Yoda1 is a specific activator of PIEZO1, and *Plasmodium* species do not contain PIEZO orthologues, Yoda1 could have a PIEZO-independent activity on the parasite itself. To explore this possibility, uninfected RBCs were first pre-treated with Yoda1 or with DMSO for 10 min and then extensively washed to remove Yoda1. Washed RBCs were then mixed with purified segmenter stage infected RBCs which were close to egress, and incubated overnight before determining parasitemia 18 hours later. Again, a strong reduction in parasitemia was observed, although shifted to a higher concentration range with an IC50 of about 3 μM (****Figure 1B****). The effect was independent of a haemolytic activity of Yoda1 that was only observed at higher concentration (10 μM) and for prolonged incubation times (24 h) (****Supplemental figure 1A****).

Thus, the reduction in parasitemia is due to an effect of Yoda1 on the erythrocyte and the effect is reversible for concentrations lower than 1 μM, without involving a haemolytic activity.

### Yoda1 prevents RBC invasion by *P. falciparum* merozoites

The observed reduction in parasitemia caused by Yoda1 could be due to selective lysis of infected RBCs, reduced growth and development of parasites, interference with parasite egress from the infected RBC, reduction of RBC invasion, or to a combination of these effects. We observed no reduction in parasitemia when synchronised ring stage parasites were incubated with Yoda1 for 24 h (****Figure 2A****), arguing again against Yoda1-induced selective lysis. Moreover, progression through the different stages of parasite maturation was not affected in the presence of 5 μM Yoda1 (****Figure 2B****). The same number of daughter cells were formed in the presence or absence of 5 μM Yoda1 (****Figure 2C****). Altogether, these results indicate that intracellular parasite development is not affected by Yoda1.

**Figure 2.**
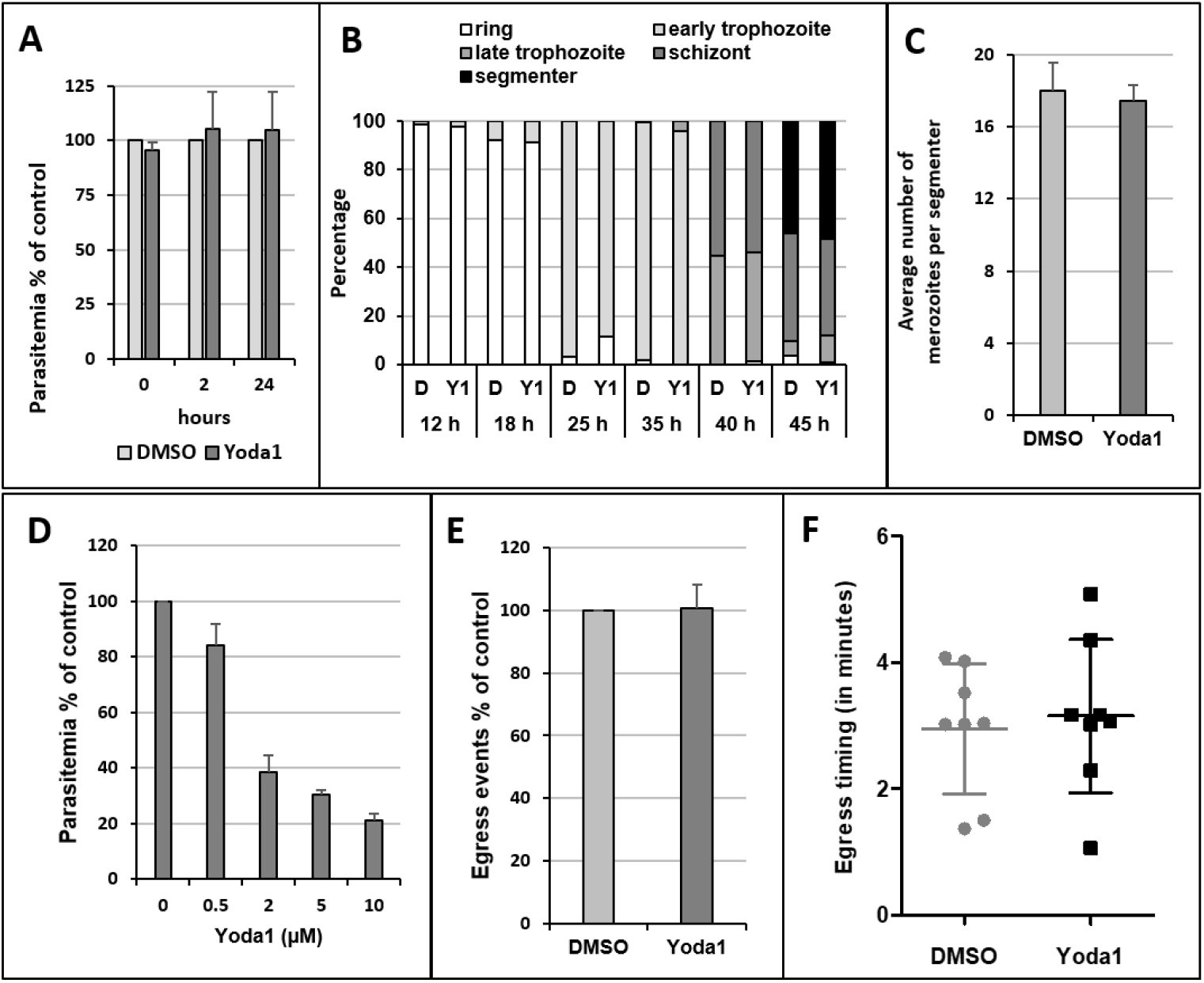
Yoda1 does not affect intra-erythrocytic growth and development. (**A**) Yoda1 (5 μM) was added to ring stage parasites (0h to 4h post invasion) and parasitemia determined after 2h and 24 h incubation (n=3). (**B**) The progression through the intra-erythrocytic development in the presence of 5 μM Yoda1 was followed by Diff-Quick-stained thin smears. At 40 h cultures were treated with PKG inhibitor C2 to block egress. Shown are the results of one representative experiment of 3. (**C**) The number of merozoites formed per segmenter was counted on stained thin smears after 45 h culture in the presence of 5 μM Yoda1. Egress had been blocked by C2 treatment (n=2). (**D**) Percoll-purified late-stage parasites were used as starting material for cultures in the presence of Yoda1 for 18 h (n=3). Kruskal-Wallis test p<0.01. (**E**) Relative number and (**F**) timing of egress events were determined from video microscopy analysis. Purified segmenter-stage parasites had been blocked before egress with C2. Upon removal of inhibitor by a rapid wash, parasites were added to complete medium containing 5 μM Yoda1 or DMSO vehicle and monitored in parallel allowing comparison of the total number of egress events and the relative time to first egress (after start of the video acquisition) between the two conditions in two independent experiments. **A** to **F**: mean + SD.

When we purified synchronized late-stage parasite-infected RBCs and incubated them with fresh RBCs in the presence of Yoda1 for 18 h and measured parasitemia by blood smear or FACS, we observed a similar dose-dependent inhibition, indicating that either egress or invasion was affected (****Figure 2D****). We observed parasite egress by live microscopy in the presence of 5 μM Yoda1. Highly synchronised late-stage parasite-infected RBCs were purified and allowed to mature to the segmenter stage in the presence of the protein kinase G (PKG) inhibitor C2; this treatment reversibly blocks parasite egress from the host RBC. The block of egress was then lifted in the presence of Yoda1 or of vehicle and both samples were observed in parallel by live microscopy. Counting the number of egress events (****Figure 2E****) and determining the relative time of the first egress event revealed that there was no difference between the two samples (****Figure 2F****).

Thus, Yoda1 inhibits invasion of human RBCs by *P. falciparum*, but does not interfere with parasite growth or egress.

#### Yoda1 protection against *P. falciparum* is independent of erythrocyte echinocytosis

Taking advantage of video microscopy, we examined the change in cell shape of RBCs induced by Yoda1. Within 30 seconds after Yoda1 addition human RBCs became echinocytic, as previously reported for mouse RBCs (****Cahalan et al., 2015****) (****Video 1****). We also followed the development of RBC morphology over more prolonged periods of time (up to three hours) in the presence of Yoda1 (****Figure 3A, Video 2****). Video-microscopy revealed that Yoda1 treated RBCs changed their shape over time from echinocytes to rounded cells (discocytes or stomatocytes); over 80% of cells had spontaneously changed after 90 min incubation at 37°C (****Figure 3A****).

**Figure 3.**
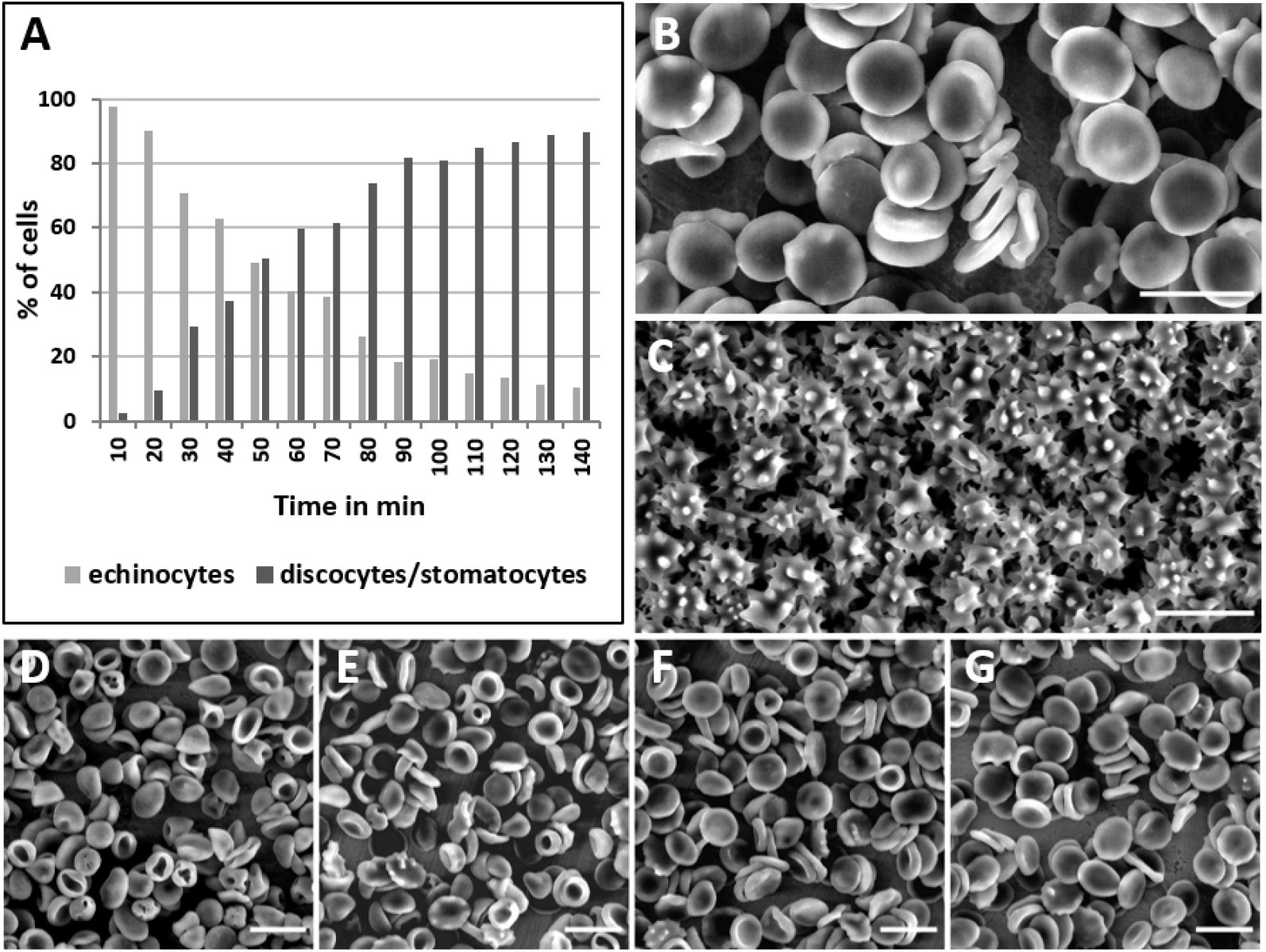
RBC shape in the presence of Yoda1 and upon washout. **(A)** Graphical representation of cell shape changes over time as observed by live microscopy in complete medium in the presence of 10 μM Yoda1 (see ****Video 2****). **(B-G)** Scanning electron microscopy images of RBCs. (**B**) DMSO mock treated, (**C**) in the presence of 5 μM Yoda1, **(D-G)** after treatment with 5 μM Yoda1 for 10 min, followed by washes and incubation in complete medium at 37°C for 30 min (**D**), 2 h (**E**), 6 h (**F**), and 24 h (**G**).

Scanning electron microscopy (SEM) images of RBCs in the presence of Yoda1 further examined in detail the changes in cell shape (****Figure 3B, 3C****). All RBCs had changed to echinocytes that were characteristic in having many tiny spikes. We also analysed RBCs that had first been treated for 10 min with 5 μM Yoda1, and then washed twice and incubated for 30 min, 2 h, 6h, or 24 h before fixation (****Figure 3D-G****). These RBCs showed no more spikes and resembled discocytes and stomatocytes.

Importantly, RBCs pre-treated for 10 minutes with 5 μM Yoda1 and then washed for 30 minutes or longer prior incubation with the parasites still showed blunted parasitemia. Thus, protection of Yoda1 against *P. falciparum* still occurs when RBCs have recovered their discoid shape.

### Protection against *P. falciparum* infection is not mediated by RBC dehydration

Previous findings indicate that dehydrated RBCs are less susceptible to *Plasmodium* infection (***Tiffert et al., 2005***). Normal RBCs that had been experimentally dehydrated or RBCs from donors with known hereditary conditions that result in dehydrated RBCs (including sickle cell disease) are greatly protected against *P. falciparum* infection. In the same line, Yoda1, as well as GOF *PIEZO1* mutations (slowing down inactivation) cause mouse RBCs dehydration in a defined minimal medium containing 2 mM Ca^2+^, as revealed by a leftward shift of the osmotic fragility curve (***Cahalan et al., 2015; Ma et al., 2018; Rapetti-Mauss et al., 2017***). Here, we investigated whether or not RBC dehydration contributes to the antimalarial activity of Yoda1 in our standard parasite culture conditions.

Parasites in human RBCs were cultured in RPMI 1640 that contains 0.42 mM Ca^2+^ and supplemented with 0.5% albumax, a standard medium that guaranties parasite viability and growth (****Schuster, 2002****). Strikingly, under this specific culture condition, Yoda1 shifted the osmotic fragility curves to the right, indicating that human RBCs become slightly overhydrated (instead of dehydrated) in the presence of Yoda1 (****Figure 4A****). However, the Ca^2+^ ionophore A23187 still caused dehydration (i.e. leftward shift in the osmotic fragility curve) in this culture medium. Similar results were obtained with mouse RBCs cultured in RPMI medium (****Figure 4B****). However and in agreement with previous findings (***Cahalan et al., 2015***), with a defined minimal medium containing 2 mM Ca^2+^, we observed a left shift of the osmotic fragility curves in the presence of Yoda1, reflecting human RBC dehydration (***Figure 4C***). A key difference between both culture media is the calcium concentration: 2 mM in the minimal medium versus 0.42 mM in RPMI. Next, we evaluated the calcium dependency of Yoda1-induced dehydration of human RBCs using a defined minimum medium. Notably, Yoda1 induced dehydration of RBCs only at 2 mM Ca^2+^ concentration (****Figure 4D, 4E, 4F****).

**Figure 4.**
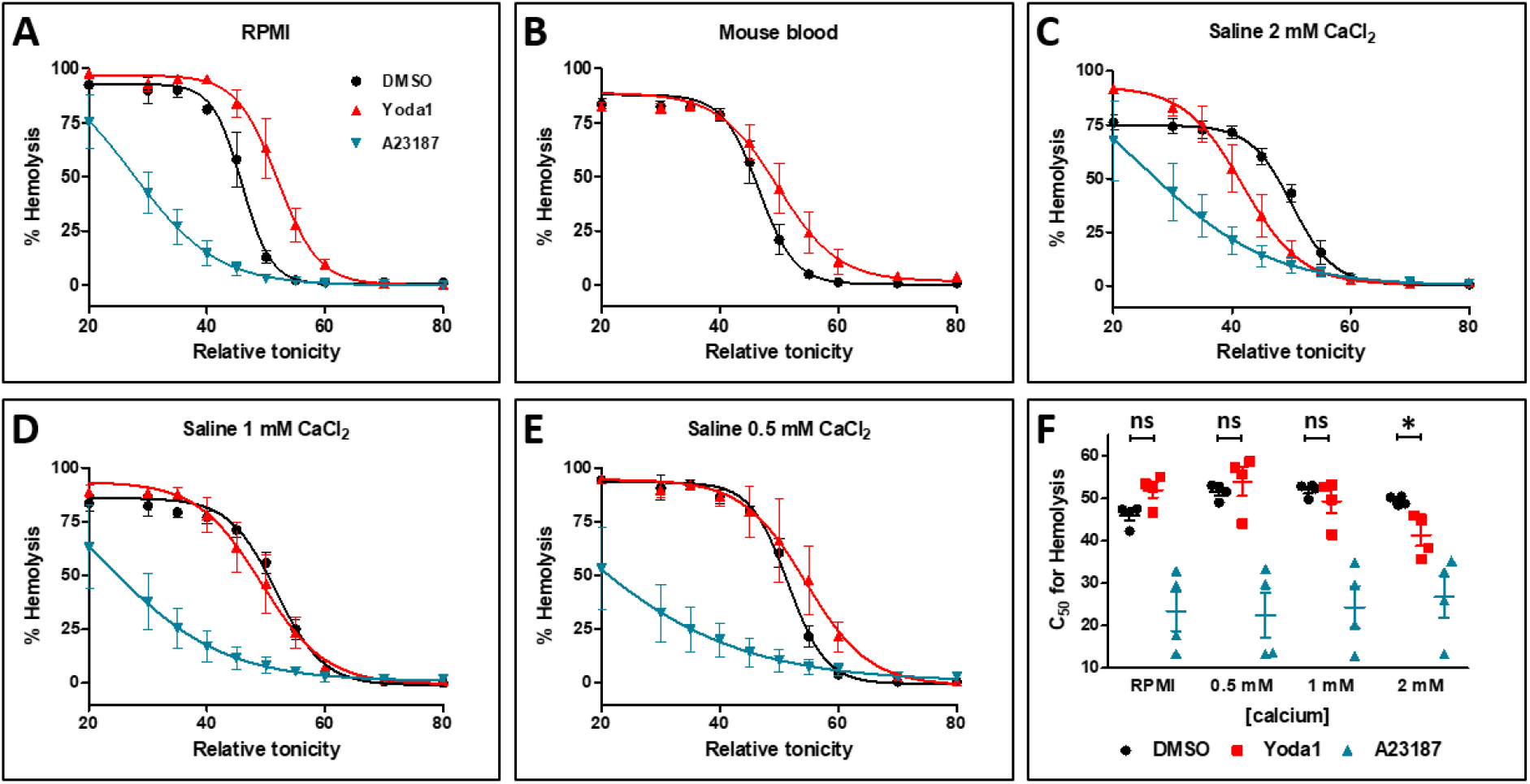
The hydration status of Yoda1 treated human RBCs. Osmotic fragility assays were performed on RBCs of different donors that had been treated for 30 min at 37°C in the indicated media with 5 μM Yoda1, 2 μM A23187, or DMSO vehicle (±SD, n=4). (**A**) RPMI, (**B**) mouse blood in RPMI, (**C-F**) minimal medium with varying calcium concentrations: 2 mM (**C**), 1 mM (**D**) and 0.5 mM (**E**). (**F**) Comparison of the LC50 values ±SEM calculated from the graphs in A, C, D, and E. Statistical analysis was done using the non-parametric Mann Whitney test (n=4), *p < 0.05. In addition, all comparisons of DMSO and A23187 samples showed p < 0.05.

Thus, in RPMI medium, the antimalarial effect of Yoda1 occurs independently of RBCs dehydration.

### Protection of Yoda1 against *P. falciparum* is independent of an altered intracellular Na^+^/K^+^ balance

PIEZO1 opening alters RBC’s intracellular cationic homeostasis (Cahalan et al., 2015; Rapetti-Mauss et al., 2017). We investigated whether or not changes in intracellular Na^+^ and K^+^ concentrations might contribute to the antimalarial properties of Yoda1. Human RBC in RPMI (i.e. the medium used to grow parasites) were treated for 10 min with Yoda1 (1 μM and 5 μM). Intracellular Na^+^ and K^+^ concentrations were determined by flame spectroscopy both immediately after the Yoda1 treatment and after an extensive wash for 24h (Figure 5A). A dose-dependent reduction in K^+^ content was readily observed with a concomitant increase in Na^+^ content, persisting after a 24 hour washout when tested at a concentration of 5 μM.

**Figure 5.**
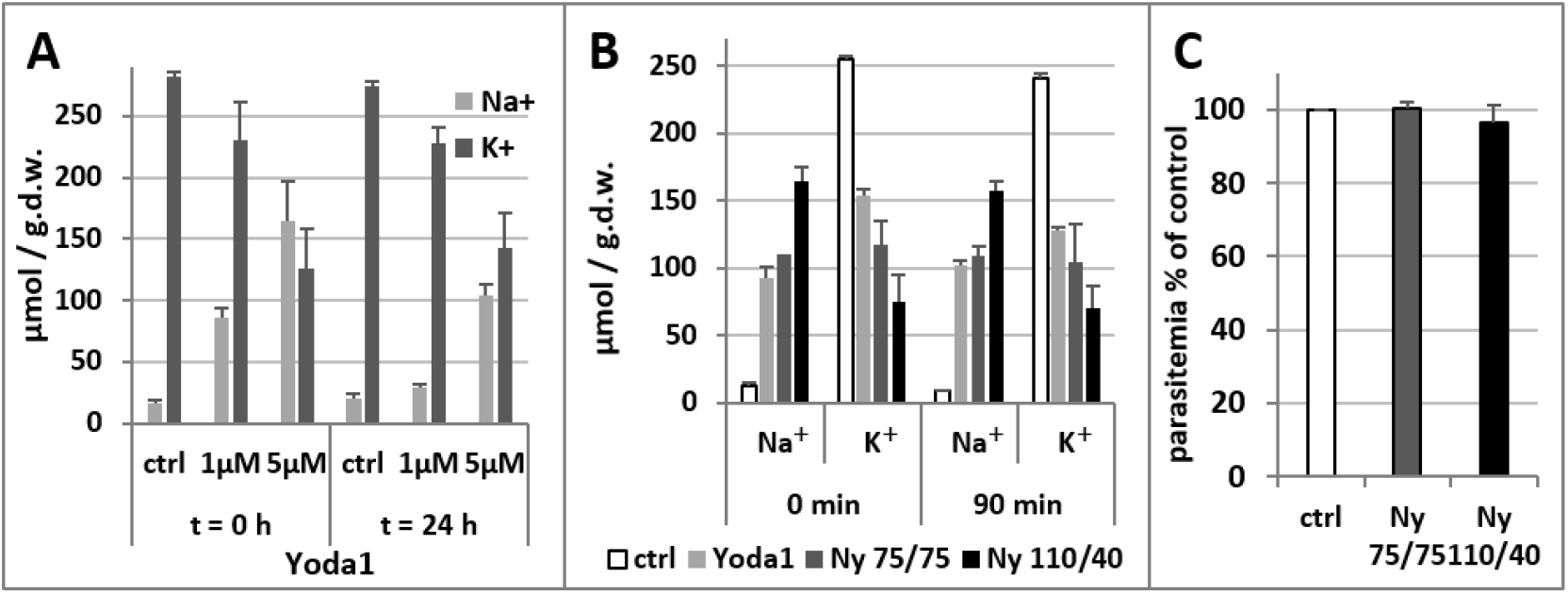
Intracellular Na^+^/K^+^ balance in human RBCs upon Yoda1 treatment and effect on *P. falciparum* infection. (**A**) Na^+^ and K^+^ content was determined upon standard 10 min treatment with 1 μM or 5 μM Yoda1 either directly after treatment and washes or after further 24h culture in complete medium at 37°C. (**B**) Treatment of RBCs with the ionophore nystatin in saline solution containing either 75 mM Na^+^ and 75 mM K^+^ (Ny 75/75) or 110 mM Na^+^ and 40 mM K^+^ (Ny 110/40). The resulting intracellular Na^+^ and K^+^ concentrations were determined directly after washing and after a further incubation for 90 min at 37°C in RPMI. (**C**) RBCs treated with nystatin in the 2 saline solutions were used for used for standard invasion assays with purified highly synchronous late-stage parasites. Results are expressed as percentage of untreated RBC controls (n=3). All graphs represent mean + SD.

Next, we explored whether or not altered intracellular Na^+^/K^+^ balance contributes to the antimalarial effect of Yoda1. We experimentally modified the intracellular Na^+^ and K^+^ content by pre-treatment of RBCs in saline solutions with varying Na^+^/K^+^ concentrations in the presence of the ionophore nystatin (****Figure 5B****). In the presence of 75mM Na^+^ / 75 mM K^+^ and nystatin, Na^+^ was elevated while K^+^ was decreased, similar to the Yoda1 condition. This effect was enhanced with the 110 mM Na^+^ / 40 mM K^+^ and nystatin solution. Changes in intracellular cationic concentration persisted after a washout of 90 minutes in RPMI (****Figure 5B****), i. e. around the time when RBCs would be invaded when cultured in the presence of purified parasites. Importantly, *P. falciparum* infection was not affected by the nystatin treatment with these modified ionic solutions (****Figure 5C****).

Altogether these findings indicate that the antimalarial activity of Yoda1 is not mimicked by experimental conditions similarly affecting the intracellular Na^+^/K^+^ balance.

### Antiplasmodial properties of Jedi1 and Jedi2

The Jedi1 and Jedi2 small molecules are chemically unrelated to Yoda1, and both activate PIEZO1 in the millimolar range (****Wang et al., 2018****). Interestingly, both Jedi1 and Jedi2 reduced *P. falciparum* parasitemia, similarly to Yoda1 (****Figure 6A****, drug on). The inhibitory effect of Jedi1 and Jedi2 on parasite invasion was readily reversible. Washing out Jedi1 or Jedi2 prior to infection by parasites completely re-established their competence for parasite invasion (***Figure 6A***, drug off). Thus, the antimalarial activity of PIEZO1 activators is not due to a deleterious effect of the compounds (that would be irreversible) on RBCs viability and protection against infection is not restricted to Yoda1. Similarly to Yoda1, in the presence of both Jedi1 and Jedi2, RBCs rapidly became echinocytes upon addition of the PIEZO1 activators (****Figure 6B****), in the absence of any haemolytic activity (****Supplemental figure 1B****). Notably, treatment with Jedi1 or Jedi2, also failed to cause RBC dehydration when cultured in RPMI (****Figure 6C****).

**Figure 6.**
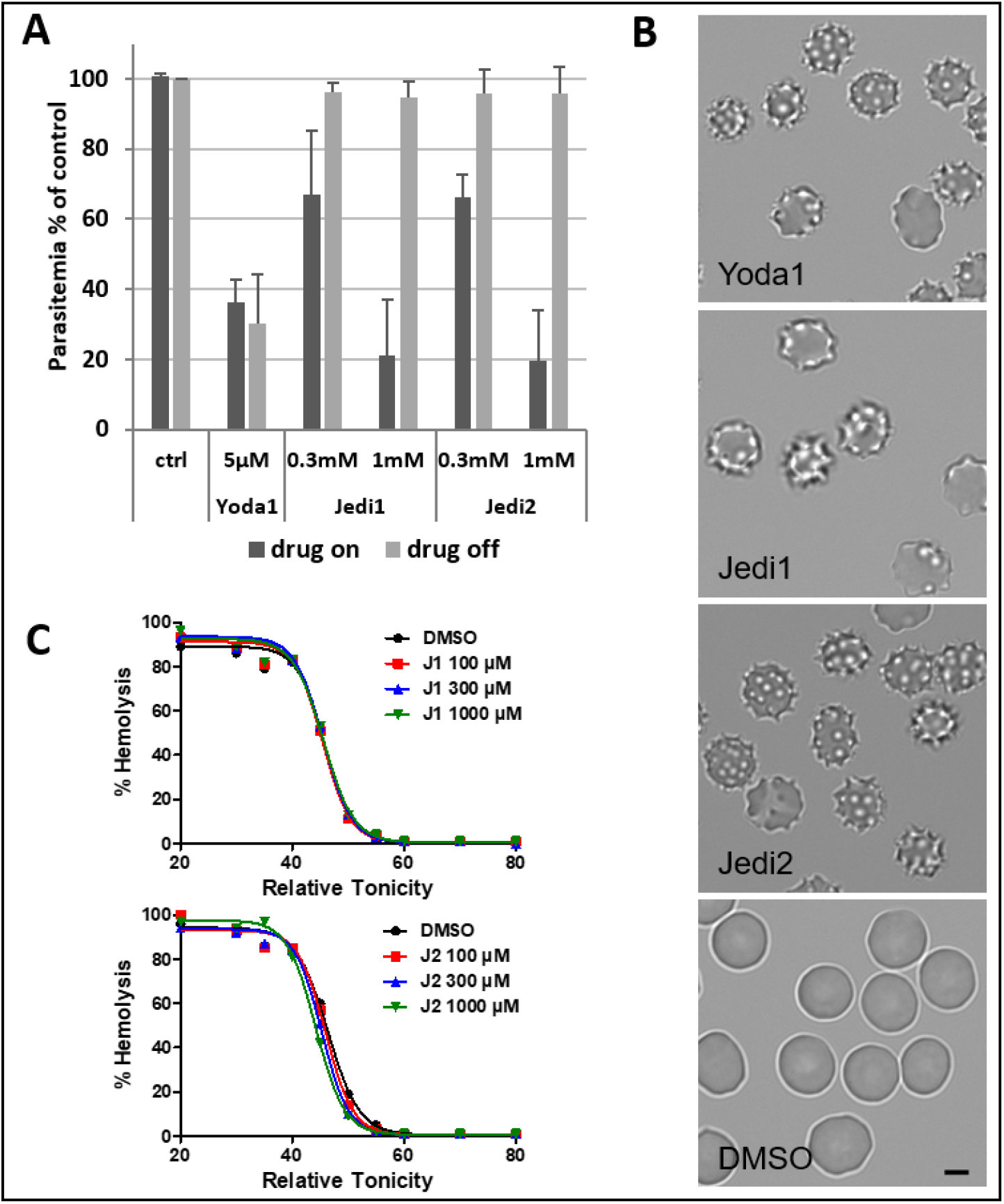
PIEZO1 activation in RBCs by Jedi1 and Jedi2. (**A**) The effect on parasitemia was observed either in the continuous presence of drug (drug on) in complete medium, or after a pre-treatment for 10 min in RPMI followed by extensive washes (drug off) before adding synchronized schizont infected RBCs for culture in complete medium. Parasitemia was determined after incubation for 18 h and expressed as percentage of mock-treated controls (n=3, mean + SD). Kruskal-Wallis test for drug on condition of Jedi1 and Jedi2 p<0.05. (**B**) Cell shape of the cells at 4 min after addition of compounds (5 μM Yoda1, 1 mM Jedi1 and Jedi2). Scale bar = 5 μm. (**C**) Osmotic fragility assays with RBCs treated with increasing concentrations of Jedi1 (top) and Jedi2 (bottom). One representative experiment (of 3) is shown.

Altogether, these findings indicate that the PIEZO1 activators Jedi1 and Jedi2, chemically unrelated to Yoda1, also prevent *in vitro* parasitemia.

### Yoda1 impairs the interaction of the parasite with the RBC membrane at multiple levels

Next, we explored the initial cellular events occurring upon invasion of control and Yoda1-treated RBCs by *P. falciparum* merozoites using direct observation by video microscopy. Highly synchronized purified segmenter stage parasites were first blocked before egress by treatment with C2. Upon washing off the inhibitor, the infected RBCs were mixed with RBCs that had been pre-treated with 5 μM Yoda1 or with vehicle only and then washed. From the analysis of the videos, we quantified the following events that occur in chronological order: a) the number of RBCs that came close to and likely came in contact with released merozoites; b) RBCs that deformed upon merozoite contact; c) invasion events; and d) echinocyte formation of RBCs as a consequence of completed parasite invasion (****Figure 7A, 7B, Video 3, Video 4****). When we expressed these data as a percentage of the number of initial RBC/merozoite contacts, we observed a major reduction in the number of early RBC deformation, merozoite invasion and echinocyte formation.

**Figure 7.**
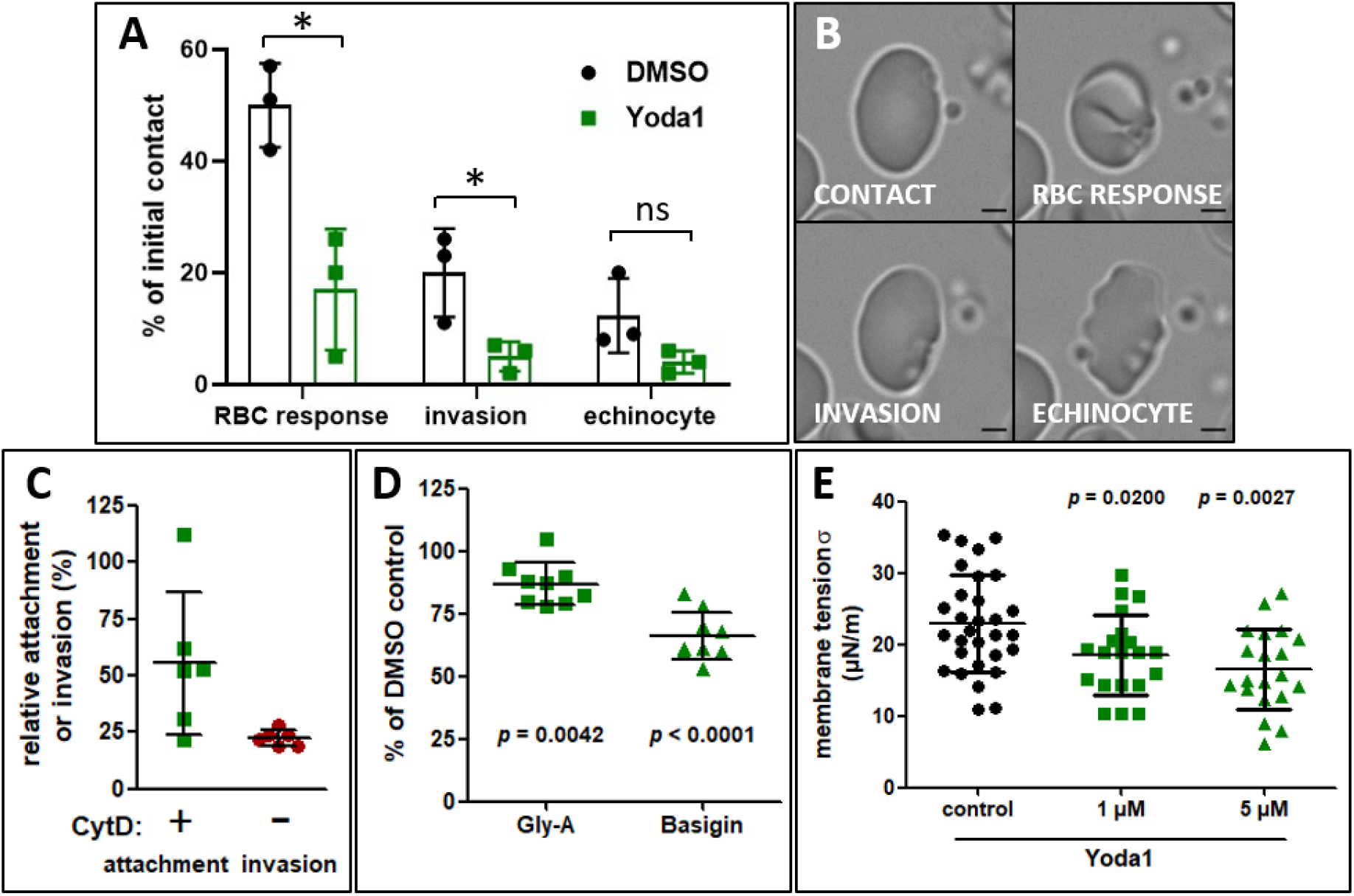
Effect of Yoda1 on the RBC and merozoite attachment and invasion. (**A**) Quantification of invasion events in the presence and absence of Yoda1. Highly synchronous purified parasites were mixed with RBC preparations that had been pre-treated for 10 min with 5 μM Yoda1 or DMSO 5 hours prior to the experiment. Events subsequent to egress were observed by live microscopy and classified. Data are expressed with respect to the observed initial possible contact of merozoites with RBCs. The results of three independent experiments are shown. * = p<0.05 unpaired two-tailed t-test. Ns = not significant (**B**) Still images of one RBC (DMSO condition, taken from video 3) representing the events that were quantified: merozoite contact, RBC response by deformation, invasion, and echinocyte formation. Scale bar 2 μM. Supplementary videos 4 represents an example of the poor reponse of Yoda1-pretreated RBCs upon contact with merozoites. (**C**) Attachment of merozoites to RBCs pre-treated with 5μM Yoda1 was quantified in the presence of 1 μM CytD and expressed as percentage of mock-treated control RBCs. In the absence of CytD we observed the usual reduction in parasitemia. Six independent experiments were performed. p<0.0001 for both conditions, Two-way ANOVA. (**D**) Quantification of the RBC surface markers GlycophorinA (Gly-A) and basigin after 5 μM Yoda1 treatment. Median fluorescence was expressed as percentage of mock-treated control RBCs. Blood of eight different blood donors was used. p-values were calculated by two-way ANOVA. (**E**) Measurement of membrane tension upon the addidtion of different concentrations of Yoda1. The tension σ was calculated from experiments presented in Supplemental Figure 2. n = 20 for Yoda1-treated cells and n = 30 for controls. p-values were calculated by Mann Whitney test. Graphs A, C, D, E show the mean ± SD.

We then determined whether or not the upstream event of merozoite attachment to RBCs would also be modified by Yoda1 treatment. We took advantage of the small molecule inhibitor of actin polymerisation, cytochalasin D (CytD) to allow attachment of merozoites to RBCs, while preventing parasite invasion (****Miller et al., 1979****) (****Paul et al., 2015****). In the presence or absence of cytD, pre-treatment of RBCs with Yoda1 followed by washes and recovery for 5 h resulted in a significant decrease in the attachment of the parasite (****Figure 7C****). Treatment with heparin that blocks adhesive events likely through interaction with the parasite invasion ligand MSP1 (***Boyle et al., 2010; Weiss et al., 2015***), completely abolished attachment and invasion and was used as baseline controls.

The observed reduction in attachment could be due to modifications of the molecular interactions between parasite ligands and RBC surface receptor proteins and/or to biophysical changes of the RBC membrane as a consequence of Yoda1 treatment. To identify their possible contributions, we first quantified by FACS analysis the surface exposure of RBC surface proteins in Yoda1- and mock treated RBCs. Glycophorin A has been identified as the receptor of EBA-175 (****Orlandi et al., 1992****) and basigin as the receptor of the RH5 complex (***Baum et al., 2009; Crosnier et al., 2011***). The median fluorescence for both surface proteins was significantly reduced for Yoda1-treated cells with respect to DMSO-treated RBCs (****Figure 7D****).

We finally analysed whether or not membrane tension was altered in the presence of Yoda1. RBC membrane fluctuations were measured in the presence of Yoda1 and the surface membrane tension σ was determined (Supplemental Figure X). The membrane tension σ significantly decreased dose dependently (****Figure 7E****).

## Discussion

Recent work demonstrates that polymorphisms in the *PIEZO1* gene causing a channel GOF, with a slower inactivation (i.e. increased channel opening), are associated with a protective effect against severe malaria both in humans and in a xerocytosis mouse model (***Ma et al., 2018; Nguetse et al., 2020***). However, the PIEZO1-dependent mechanisms at play conferring the resistance to severe malaria remains debated and was previously suggested to be caused by RBC dehydration (***Ma et al., 2018***) or to impaired export of *P. falciparum* virulence protein PfEMP-1 (***Ma et al., 2018; Nguetse et al., 2020***). Our study focuses on the *in vitro* pharmacological activation of PIEZO1 in human RBCs and its impact on invasion, growth and development of the malaria parasite *P. falciparum*.

We show that Yoda1, a potent PIEZO1 activator (****Syeda et al., 2015****), has a strong inhibitory effect on parasitemia in the low micromolar range, reminiscent of the observed reduced multiplication rates in mice carrying *PIEZO1* GOF mutations (****Ma et al., 2018****). Yoda1 does not affect parasite growth or development within RBCs, nor parasite egress from the infected RBCs. However, invasion of RBCs by *Plasmodium* merozoites is potently inhibited by Yoda1. Decrease in parasitemia is mediated by an effect of Yoda1 on RBCs, but not on the parasite itself, since inhibition of invasion is also observed when RBCs had been pre-treated with Yoda1 and washed before the addition of parasites. The chemically unrelated PIEZO1 activators Jedi1 and Jedi2, similarly inhibit *P. falciparum* parasitemia, further indicating a PIEZO1-dependent mechanism. Yoda1 treated RBCs rapidly become echinocytes and recover their cell shape after prolonged incubation or washout, while the anti-malaria effect persists. Moreover, Yoda1 induces a major imbalance in intracellular Na^+^ and K^+^ content, although it is not at the origin of the protection against malaria. Instead, we provide evidence that Yoda1 inhibits merozoite attachment, in parallel with a reduced surface exposure of *Plasmodium* receptors. Altogether, our in vitro findings further support a role for PIEZO1 in the host RBC contributing to the resistance against severe malaria in vivo conferred by a GOF mutation in PIEZO1 (****Ma et al., 2018****)

Previous studies demonstrated that GOF mutations in *PIEZO1* cause xerocytosis, a mild hemolytic anaemia (***Albuisson et al., 2013; Andolfo et al., 2013; Bae et al., 2013; Zarychanski et al., 2012***). RBC dehydration was shown in xerocytosis patients (****Andolfo et al., 2013****), in asymptomous human carriers of a *PIEZO1* E756 del polymorphism (****Ma et al., 2018****), as well as in an engineered *PIEZO1* mouse model of xerocytosis (***Cahalan et al., 2015; Ma et al., 2018***). RBC dehydration arises through PIEZO1 mediated calcium influx leading to activation of the calcium-dependent K^+^ channel Gardos and efflux of K^+^ ions accompanied by water loss *via* aquaporins (****Cahalan et al., 2015****). However, our data revealed that under standard *P. falciparum in vitro* culture conditions (in RPMI), RBCs are not dehydrated upon treatment with the PIEZO1 activators Yoda1, Jedi1 or Jedi2. Lack of dehydration in this specific culture medium is due to the low extracellular calcium concentration used (RPMI only contains 0.42 mM calcium). Thus, the observed protection mediated by PIEZO1 activators against malaria infection (at least in the RPMI medium) does not depend on RBCs dehydration. Nevertheless, an additional protective effect of RBCs dehydration against malaria was previously demonstrated in various clinical conditions such as xerocytosis and sickle cell disease, as well in a variety of experimental conditions (****Tiffert et al., 2005****). Thus, PIEZO1 opening in RBCs is anticipated to confer protection against malaria by two independent mechanisms, one involving RBCs dehydration (***Ma et al., 2018; Tiffert et al., 2005***) and the other one independent of RBCs volume regulation (****Nguetse et al., 2020****)(and the present report).

Invasion of RBCs by *P. falciparum* merozoites is a complex process that has mainly been considered from the point of view of the invading parasite (reviewed in (***Cowman et al., 2017; Weiss et al., 2016***). The sequential events for successful invasion are: 1) attachment of the merozoite to the RBC by binding to specific RBC surface proteins triggering waves of RBC deformation; 2) reorientation of the parasite resulting in the apical pole facing the RBC; 3) discharge of apical organelles (micronemes and rhoptries) leading to the formation of a moving junction; 4) invasion by active movement of the parasite using the moving junction as a fixation point for the activity of a parasite actin-myosin motor, leading *in fine* to the formation of a parasitophorous vacuole in which the parasite will develop during its intraerythrocytic growth (***Gilson and Crabb, 2009; Riglar et al., 2011***).

The deformation of the RBC during the initial interaction with the merozoite is anticipated to activate PIEZO1. However, no calcium signal in the RBC was reported during the pre-invasion stages (attachment and reorientation) when RBCs maximally deform upon initial merozoite contact, suggesting that PIEZO1 activation is unlikely to occur at this stage (***Introini et al., 2018; Weiss et al., 2015***). Binding of merozoites to human RBCs triggers important biophysical changes that will contribute to facilitate invasion (****Koch et al., 2017****). Modelling has suggested that wrapping of the RBC cell membrane followed by reorganisation of the underlying cytoskeleton could account for the energetic steps required for the invasion process (****Dasgupta et al., 2014****). Upon contact with the parasite, a signalling pathway through a phosphorylation cascade causes altered viscoelastic properties of the host membrane (****Sisquella et al., 2017****). A possibility is that these early events may prevent PIEZO1 opening at the very beginning of the invasion process (despite major membrane deformation), thus explaining the absence of a calcium signal at this stage where membrane movements are remarkably large (***Introini et al., 2018; Weiss et al., 2015***). Another question is whether or not the intraerythrocytic parasite may also directly influence PIEZO1 activity within the host cell. Differential phosphorylation of PIEZO1 has been documented in parasitized RBCs (****Bouyer et al., 2016****). Moreover, in a quantitative phosphoproteomic analysis of the erythrocyte during the short phase of invasion, PIEZO1 was one of the few identified proteins showing increased phosphorylation upon merozoite attachment prior to invasion (****Zuccala et al., 2016****). However, the biological significance of PIEZO1 phosphorylation in infected RBCs is unclear at this stage.

Multiple questions remain open to better understand how the pharmacological activation of Piezo1 confers a potent protection against malaria infection. Recent structural findings indicate that PIEZO1 negatively curves the membrane in the closed state (***Guo and MacKinnon, 2017; Lin et al., 2019; Saotome et al., 2018; Zhao et al., 2018***). Upon activation in response to mechanical stimulation, the channel reversibly flattens (****Lin et al., 2019****). How the effect of PIEZO1 on RBC membrane curvature may possibly influence merozoite attachment (although membrane flattening is anticipated to enhance access of parasite to RBC surface receptor proteins) and deformation of the RBC membrane upon initial contact with the parasite is an intriguing question that deserves to be addressed. Does the decrease in RBC membrane tension observed in the presence of Yoda1 contribute to the reduced parasite interaction and invasion? Moreover, whether or not membrane depolarization (although the resting membrane potential of human RBC is depolarized in the range of −10 mV) and/or the increase in intracellular calcium (although blunted in our standard RPMI medium) are also involved in the mechanism of protection is unknown at this stage. In the same line, PIEZO1 opening in RBCs is anticipated to activate several downstream biochemical cascades (presumably calcium-dependent) impacting various structural components of RBCs, including the cytoskeleton (****Li et al., 2014****), as well as the lipid bilayer composition by influencing the cellular lipidome (****Buyan et al., 2020****). How these changes may contribute to the protection against malaria infection also need to be investigated in future studies.

The *PIEZO1* E756del polymorphism is highly enriched in people of African origin (****Ma et al., 2018****). Remarkably, this variant strongly associates with significant protection against severe malaria, although these heterozygote individuals have no detectable clinical symptoms caused by this deletion (****Nguetse et al., 2020****). The mechanisms implicated in the protection conferred by the *PIEZO1* E756del against severe malaria remain disputed (***Ma et al., 2018; Nguetse et al., 2020***). Our *in vitro* findings now indicate that PIEZO1 pharmacological activation in human RBCs is sufficient to prevent invasion by *P. falciparum,* associated with a reduction in RBC surface receptors of *Plasmodium*, merozoite attachment and subsequent RBC deformation. In conclusion, it is likely that a combination of effects involving inhibition of invasion (the present report), decreased expression of the cytoadherence molecule PfEMP-1 (****Nguetse et al., 2020****), RBC dehydration (***Ma et al., 2018; Tiffert et al., 2005***), and modulation of the immune system (****Ma et al., 2018****) mediates the PIEZO1-dependent protection against severe malaria.

## Materials and methods

### Chemicals

The following chemicals and their stock concentrations in DMSO (in square brackets) were used in this study: Yoda1 (Sigma-Aldrich)[20 mM], A23187 (Sigma-Aldrich)[2 mM], Jedi1 (MolPort, Riga, Latvia)[200 mM], Jedi2 (MolPort)[200 mM], Nystatin (Sigma-Aldrich)[40 mg/ml]. The PKG inhibitor C2 was a generous gift of Oliver Billker [3 mM].

### Biological material

Cell cultures of *P. falciparum* 3D7 strain (MRA-102 from MR4) were performed as previously described (****Trager and Jensen, 1976****) in human 0^+^ or A^+^ erythrocytes and complete medium, RPMI 1640 medium supplemented with 25 mM HEPES and 2 mM L-glutamine (Gibco), 0.5% Albumax I (Gibco), 15 μg/ml hypoxanthine (Sigma-Aldrich) and 40 μg/ml gentamycin (Gibco) at 37°C under a tri-gaseous mixture of 5% CO2, 5% O2 and 90% N2. Culture conditions were generally static for culture maintenance and shaking for experiments to obtain higher multiplication rates and single infected RBCs. Parasites were synchronized using VarioMACS magnetic cell separator (CS columns, Miltenyi Biotec, Paris, France) (****Ahn et al., 2008****) or centrifugation on a 63% Percoll (GE Healthcare) cushion (****Saul et al., 1982****) to purify late-stage parasites, addition of fresh RBCs, followed by a 5% sorbitol treatment 2 h to 4 h later (****Lambros and Vanderberg, 1979****). Human blood obtained as donations from anonymized individuals was provided by the local blood bank (Etablissement Français du Sang) under the approval number 21PLER2016-0103. Parasitemia was routinely monitored on thin blood smears fixed in methanol and stained with Diff-Quick™ (pH 7.2) (Dade Behring, France) or by SYBR Green staining and flow cytometry. For microscopic determination of parasitemia, at least 1000 cells were counted. Experiments were generally started with late stage parasites to eventually obtain ring stage parasitemia in the range of 1% and 10%.

### Parasitemia determination by FACS

Cultures were centrifuged and the cells fixed for at least 4 h at room temperature in 10 volumes of 4% paraformaldehyde (Electron Microscopy Sciences) in phosphate buffered saline (PBS). Preparations were then washed twice with PBS and incubated 30 min with 3.3× SYBR Green (Thermo Fisher Scientific). DNA dye was removed by a simple wash in PBS and samples analysed on a FACS Canto (BD Biosciences) using processed uninfected RBCs as negative control.

### Treatments with Yoda1, Jedi1 and Jedi2

Yoda1/Jedi1/Jedi2 treatments were performed in two different ways: either in the presence of drug by adding Yoda1 directly to the parasite cultures in complete medium (drug-on experiments) or by pre-treating uninfected RBCs, washing and infection with parasites (drug-off experiments). For the latter, RBCs in RPMI were treated for 10 min at room temperature, washed 3 times in RPMI, resuspended in complete medium and mixed with purified infected RBCs that were close to egress. To do so, late-stage parasites from synchronous cultures were purified either by passage over VarioMACS columns or by density centrifugation using a 63% Percoll/RPMI cushion. Pure infected RBCs were washed with complete medium, resuspended in complete medium with 1.5 μM of the PKG inhibitor C2, gazed and incubated at 37°C for up to 5 h. This allows parasites to continue their development to segmenter stages but blocks egress until the inhibitor is removed by washing. These parasite preparations were then mixed with either Yoda1/Jedi1/Jedi2 pre-treated RBCs or with mock-treated RBCs. To treat ring stage parasites, purified late stage parasites were mixed with RBCs, incubated for 4 h and synchronized by 5% sorbitol treatment. Cultures were allowed to stabilize for 1 h before adding Yoda1. Individual experiments were generally performed in triplicates and parasitemia determined by FACS. Experiments were repeated at least three times with blood from different donors.

### Haemolytic activity assays

Blood of different donors was obtained. Throughout these assays RPMI 1640 without phenol red (Gibco) was used. RBCs were washed twice in RPMI and suspensions at 5% haematocrit were prepared in complete medium (RPMI 0.5% albumax) and 100 μL samples were dispatched in round-bottom 96 well culture plates. Yoda1, Jedi1 and Jedi2 were prepared in complete medium at double concentration, and 100 μL added to the wells containing 100 μL RBC suspensions in triplicates (2.5% final hematocrit) and mixed by pipetting. Final concentrations for Yoda1 were 10 μM, 5 μM, 2 μM and 1 μM and for Jedi1 and Jedi2 1000 μM, 300 μM and 100 μM. Wells with DMSO were prepared for baseline determination (0.1% for Yoda1, 0.5% for Jedi). Plates were incubated in standard conditions at 37°C in 5% O2, 5% CO2. After 24h, 2 μL of saponin (15%) were added to selected wells to obtain total lysis. RBCs were pelleted by centrifugation for 3 min at 1800 g and 150 μL of supernatant transferred to flat bottom 96 well plates. Haemoglobin released into the culture supernatant was quantified using a SPARK multifunctional microplate reader (TECAN) at 540 nm. Lysis was calculated after deduction of DMSO treated samples as percentage of total saponin-induced lysis.

### Osmotic fragility assays

Osmotic fragility assays were performed as described (****Cahalan et al., 2015****) with minor modifications (****Tiffert et al., 2005****). Briefly, blood from different donors collected in citrate/dextrose was used the day of reception whenever possible. The blood was washed twice either in RPMI or in normal saline (NS, 149 mM NaCl, 2 mM HEPES, pH 7.4) depending on the following experimental conditions. One millilitre blood suspensions at 2.5% haematocrit were incubated with compounds for 30 min at 37°C in RPMI or in NS supplemented with 4 mM KCl, and variable concentrations of CaCl2 (2 mM, 1 mM, 0.5 mM). DMSO concentrations never exceeded 0.5% in all assays. Ten μl of blood suspension were added to U-bottom 96-well plates. Solutions of varying tonicity were generated by mixing NS (100%) with 2 mM HEPES, pH 7.4 (0%). 250 μl of these solutions were added to the diluted blood and incubated for 5 min at room temperature. Plates were then centrifuged at 1000 g for 5 min and 150 μl of the supernatant was transferred to flat bottom 96-well plates. Absorbance was measured using a SPARK multifunctional microplate reader (TECAN) at 415 nm. LC50 values were determined by fitting the data to 4-parameter sigmoidal dose–response curves using Prism (GraphPad).

### Video microscopy

For the observation of the cell shape, RBCs were washed and re-suspended in complete medium and settled in a poly-L lysine (PLL) coated slide chamber (Ibidi μ-slide VI). The chamber was then flushed with complete medium containing 5 μM Yoda1 and time–lapse videomicroscopy recorded with 63 x magnifications at a speed of 1 image/4 seconds. To observe the recovery of cell shape, RBCs in complete medium were treated with 10 μM Yoda1 (at this concentration, all RBCs become strong echinocytes), placed in glass bottom dishes (MatTek), and time–lapse videomicroscopy recorded with 63 x magnifications over 2 ½ h at 6 images/min.

For the observation of egress and invasion events, late-stage parasites were purified on VarioMACS columns, washed and re-suspended in complete medium and incubated for up to 5 hours in the presence of 1.5 μM PKG inhibitor C2 to block egress. For egress studies, parasites were washed twice and re-suspended in complete medium containing 5 μM Yoda1. For control samples 0.5% DMSO was used. 100 μl of each cell suspension was loaded into adjacent wells (Ibidi μ-8 well slide) and recorded in alternation. For invasion, washed parasites were mixed with RBCs that had been pre-treated with 5 μM Yoda1 or 0.5% DMSO for 10 min in RPMI, washed three times and incubated at 37°C for 5 hours. Samples were loaded in glass bottom dishes (MatTek). All events were imaged using an inverted brightfield microscope (Axioobserver, Zeiss), equipped with an incubation chamber set at 37°C and 5% CO2 and a CoolSNAP HQ2 CCD camera (Photometrics). Time-lapse experiments were performed, by automatic acquisition of designated fields using a 63× Apochromat objective (NA 1.4). Image treatment and analysis were performed using Zen software (Zeiss).

### Scanning electron microscopy

For environmental scanning electron microscopy (ESEM), samples were fixed at room temperature with 2.5% glutaraldehyde in 0.1M cacodylate buffer followed by 1% osmium tetroxide, and washed in water. A 10 μl drop of suspension was loaded on the sample carrier and imaged in a FEI Quanta200 FEG microscope in ESEM mode using the gaseous secondary electron detector. The stage was set-up at 2°C, the acceleration voltage was 15kV and the working distance 10 mm. Water was then progressively removed by cycles of decreasing pressure / injection of water, until reaching equilibrium at the dew point. The minimal final pressure in the chamber was 350 Pa. Pictures were taken with a dwell time of 6 μsec.

### Generation of RBCs with altered Na^+^/K^+^ content

To generate RBCs with a cellular Na^+^ and K^+^ content similar to the situation observed upon treatment with Yoda1, RBCs were incubated on ice in the presence of 40 μg/ml of the ionophore nystatin (Sigma-Aldrich) in a saline solution of either 75 mM NaCl / 75 mM KCl or 110 mM NaCl / 40 mM KCl in 5 mM Hepes pH 7.4 and 55 mM sucrose. Upon incubation during 20 min, cells were collected by centrifugation and washed twice for 20 min on ice with the identical solution containing 0.1% BSA to efficiently remove nystatin. The cellular Na^+^/K^+^ content was measured by spectroscopy with a Solaar spectrometer immediately after the washing and after an additional incubation of the RBCs for 90 min at 37°C in RPMI.

### Sodium and potassium measurements

Fresh venous blood was washed 3 times at room temperature in RPMI and set at 30% haematocrit. Three aliquots of 1.5 ml were treated with 1 μM or 5 μM Yoda1 or DMSO (control) for 10 min at room temperature, then rinsed 3 times in RPMI. For Na^+^ and K^+^ content measurements, 500 μl of cell suspension were taken to fill 3 nylon tubes that were centrifuged for 10 minutes at 4°C, 20,000 g at the end of Yoda1 washing (t=0) and after 24h incubation at 37°C and 5% CO2in complete medium. The pellet of RBCs was extracted. Dry weight was measured after overnight heating (80°C). Intracellular ions were extracted from dried pellets by overnight incubation at 4°C in 5 ml milliRho water (Millipore). Na^+^ and K^+^ were measured by flame spectroscopy with a Solaar spectrometer.

### RBC attachment assays

Merozoite attachment to RBCs was quantified in the presence of the inhibitor of actin polymerisation cytochalasin D (CytD) following the protocol of Paul et al (****Paul et al., 2015****). RBCs were washed twice in RPMI and treated with either 5 μM Yoda1 or DMSO vehicle for 10 min at RT. Preparations were then washed 3 times in RPMI and eventually resuspended in complete medium (RPMI, 0.5% albumax) and cultured for 4-5 h in standard conditions. Schizont- and segmenter-infected RBCs were purified from highly synchronised parasite cultures by magnetic purification on MACS columns (Milteny), effectively removing uninfected RBCs. Purified infected RBCs were cultured at 37°C for 4-5 h in complete medium in the presence of 1.5 μM C2 to allow parasite maturation while blocking egress. C2 was removed by centrifuging and resuspending cells in an equal volume of fresh complete medium. 100 μL of cell suspension were immediately added to 2ml tubes containing 100 μL of Yoda1 or DMSO pretreated RBC suspensions in the presence of 1 μM CytD (allowing attachment but inhibiting invasion), 200 μg/ml heparin (inhibiting attachment and invasion) or 0.1% DMSO vehicle (allowing invasion). Assays were done in triplicates at a final haematocrit of 2.5%. Tubes were gazed for 3 sec with 5% O2, 5% CO2 and incubated for 3 h at 37°C on a spinning wheel for mixing. Cells were fixed by adding to each tube 800 μL of a sucrose-stabilised solution (0.116 M) in PBS supplemented with 2% glutaraldehyde (***Miller et al., 1979***). The starting parasitemia (3% to 6%) was determined by fixing cells immediately after addition of iRBCs. Samples were stored at 4°C until later processing.

For quantification of attachment by FACS, 10^7^ cells (200 μL fixed suspension) were transferred to microtubes, pelleted at 300 g for 90 sec and blocked with 300 μL of a sucrose-stabilised solution (0.116 M) in PBS supplemented with 0.1 M glycine for 30 min at RT. Cells were washed twice with 500 μL PBS/sucrose and then incubated 30 min with 3.3× SYBR Green (Thermo Fisher Scientific). DNA dye was removed by a simple wash in PBS/sucrose and samples analysed on a FACS Canto (BD Biosciences) using processed uninfected RBCs as negative control. RBCs with attached merozoites (in the presence of CytD) are detected with a fluorescence signal identical to ring stage iRBCs (in the absence of CytD and heparin) while heparin treatment prevents attachment and invasion. Data were expressed as percentage of DMSO-treated control RBCs after subtraction of baseline measurements of parasitemia in the presence of heparin.

### Quantification of RBC surface markers by flow cytometry

Blood of different donors was washed twice in RPMI, treated for 10 min with 5 μM Yoda1 or 0.1% DMSO in RPMI at RT, washed three times in RPMI. Expression of the surface markers Glycophorin A (CD235a) and basigin (CD147) was monitored on 10^7^ RBCs. The fluorochrome-conjugated monoclonal antibodies GlyA-PC7 (Beckman Coulter) and basigin-APC (eBIOSCIENCES) were used at 1:100 dilution. Cells were incubated in the dark for 20 min in 50 μL PBS containing 2% fetal bovine serum (FBS) at 4°C and then washed once in the same medium at 300 g for 3 min prior to evaluation (****Kinet et al., 2007****). Analyses were performed on a FACS-Canto II cytometer (BD Biosciences). 50,000 events were recorded for each staining. Data analyses were performed using FlowJo software (Tree Star).

### Determination of membrane tension by fluctuation-based measurements

Membrane tension measurements were performed using a custom-built bright field microscope with optical tweezers based on a 1064 nm laser for back focal plane interferometry (****Allersma et al., 1998****). The optical trap was formed by two identical 63× water-immersion objectives with 1.2 numerical aperture (C-Apochromat, Zeiss). The laser power was set at 3 mW resulting in 50 μW laser beam at the backfocal plane of the objective. The backfocal plane detection was performed with a position sensitive detector (PSD, DL100-7-PCBA3 First Sensor Inc.). The voltage signal from the PSD sensor was amplified with variable gain amplifier (×10-50) to increase detection resolution. The temperature in the cell chamber was controlled with two resistor rings and Harvard Apparatus temperature controller (CL-200A Dual Channel Bipolar Controller). The rings were located in both objectives and the systems was prepared one hour before the cell chamber was placed. Once the chamber in place, the whole was maintained at 37°C for one more hour before the calibration and experiment procedure as described by Betz (****Betz et al., 2009****). Experiment chambers were build using coverslips (No 1.5, 22×64, Menzel Glaeser), the surfaces were cleaned first by sonicating for 60 min in 1 M KOH, then washed with deionized purified water 10 times and dried at 70°C for 5h. To spread poly-L-lysine (PLL) evenly, the slides were incubated in humid environment in 60 μL of 0.1 μg/mL solution of PLL per slide and covered with Parafilm™ overnight and rinsed again. The treated glass slides were protected with aluminium foil and kept at 4°C. Flow cells were built from two coverslips with Parafilm™ as a spacer and glued together by placing them on a hotplate at 50°C for few seconds, creating a 10-20 μL chamber. The top coverslips had previously been perforated using a high pressure 50 μm aluminium particle sandblaster resulting in 1 mm holes placed at both ends of the chamber. The membrane amplitude fluctuations were acquired at a frequency of 20 kHz for 5 minutes.

### Statistics and data analysis

Data analysis and plots were generated using Excel (Microsoft) and Prism (GraphPad). Osmotic fragility curves were generated by fitting the data to a 4-parameter sigmoidal dose–response curve and LC50 values were calculated. Non-parametric Mann-Whitney test was used to determine statistical significance for n>3. For dose-response experiments (Figures 1A, 1B, 2D, 6A) One-Way ANOVA and non-parametric Kruskal-Wallis test was applied.

## Acknowledgements

The authors wish to thank Loic Dumas and Viviana Claveria for assistance with experiments, Ardem Patapoutian and Shang Ma for inspiring discussions, Oliver Billker for supplying C2, Naomi Taylor for supplying RBC surface marker reagents and protocols, Catherine Braun Breton and Sharon Wein for critical reading of the manuscript. We thank the Electron Microscopy facility of the University of Montpellier (MEA) for their assistance with SEM microscopy and the imaging facility MRI, member of the national infrastructure France-BioImaging supported by the French National Research Agency (ANR-10-INBS-04, Investments for the future).

This research was supported by the Fondation Méditerranée Infection Marseille to R. Lohia, a prematuration grant of the CNRS (PREMAT317) to K. Wengelnik and D. Douguet, the Agence Nationale de la Recherche under the “Investissements d’avenir” program (ANR-16-IDEX-0006) to K. Wengelnik and R. Cerdan, a DICYT-USACH project 042031BV to R. Bernal, and LabEx Numev (Convention ANR-10-LABX-0020) to M. Abkarian. E. Honoré thanks the Fondation pour la Recherche Médicale, the Agence Nationale de la Recherche (ANR-17-CE13-0012-02, ANR-19-CE14-0029-01 and ANR-20-CE14-0032-01) and the Human Frontier Science Program for funding this work.

## Supplemental Figures and Videos

**Supplemental Figure 1:**
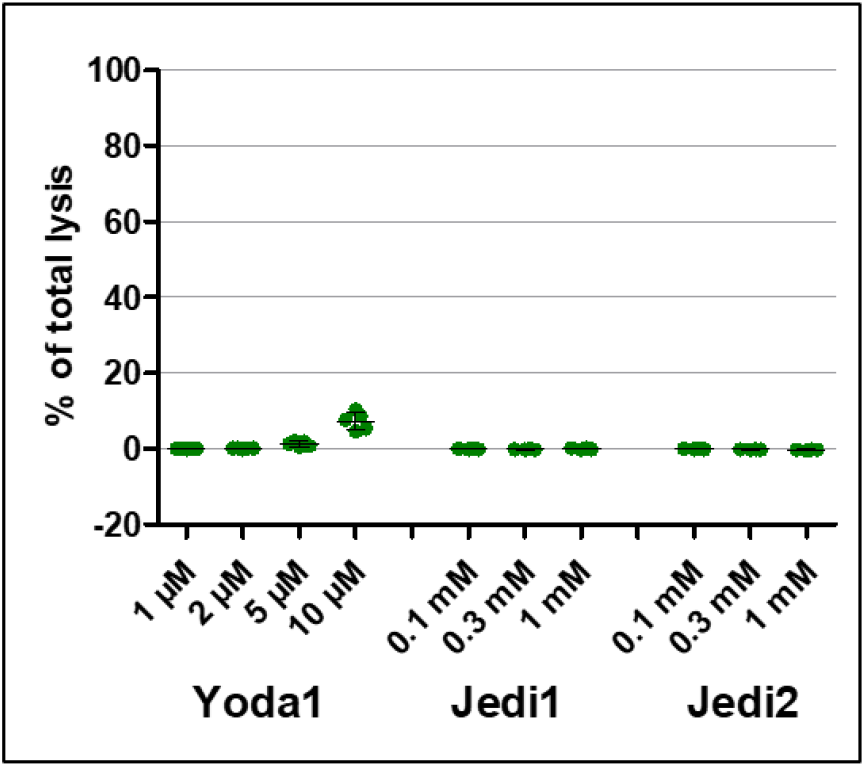
Haemolytic activities of the PIEZO1 activators Yoda1, Jedi1 and Jedi2. Blood of different donors was cultured for 24h at 2.5% haematocrit in standard culture conditions (RPMI without phenol red with 0.5% albumax) in the presence of the indicated concentrations of compounds. Haemoglobin released into the culture supernatant was quantified by measuring the absorbance at 540 nm in a TECAN spectrophotometer and expressed as percentage of total lysis induced by the addition of 0.15% saponin. Shown are mean and SD, n = 5 for Yoda1 and n = 4 for Jedi1 and Jedi2, all performed in triplicates.

**Supplemental Figure 2:**
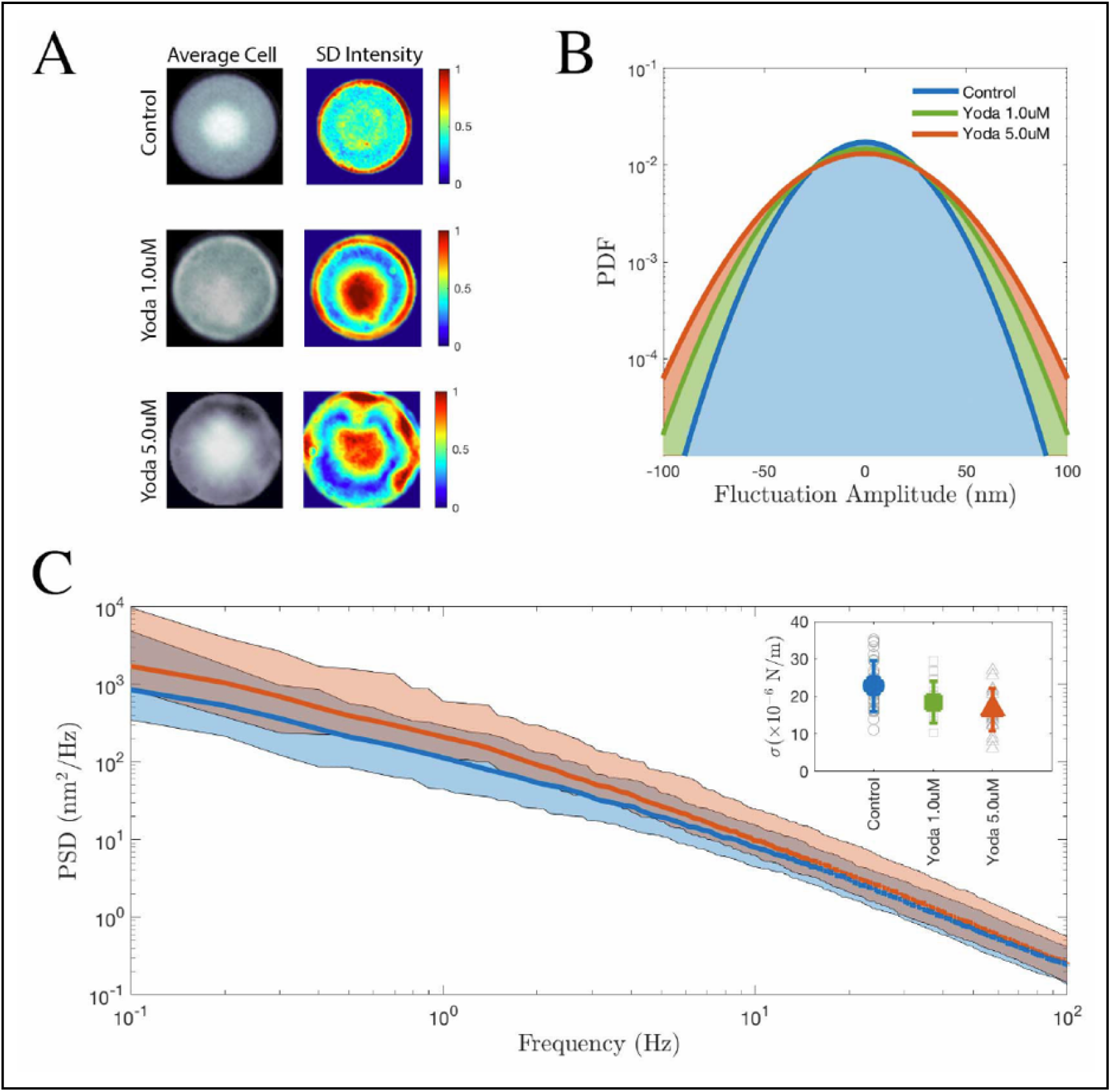
RBC membrane fluctuation in the presence of Yoda1. The cells were pre-incubated one hour in RPMI medium before the measurement started for both untreated (control) and Yoda1-treated (1 μM and 5 μM) RBCs at 37°C. **(A)** Average cell morphology (left panel) and normalized standard deviation (SD) intensity (right panel) of RBCs under the different experimental conditions. (**B)** Experimental probability density function (PDF) of the RBC membrane fluctuation amplitude displaying wider distribution at higher Yoda1 concentration indicating a reduction of membrane surface tension as a function of Yoda concentration. **(C)** Mean power spectrum density (PSD) of the membrane fluctuations (solid lines) displaying the typical variations among cells. Membrane surface tension and membrane bending modulus were extracted from fitting individual power spectrum density data. For clarity only the curves for control and 5 μM Yoda1 treatment are shown. The membrane response below 10 Hz is dominated by surface tension while above this limit the response is bending dominated. As shown in the inset, the surface membrane tension σ decreases as a function of the Yoda1 concentration (control = 22.9 (SD 6.8; n = 30), 1 μM Yoda1 = 18.5 (SD 5.6; n = 20), 5 μM Yoda1 = 16.6 (SD 5.7; n = 20) ×10^−6^ N/m) while the membrane bending modulus variations fell below 10% of the mean bending value (control: 30.4 ± 14.1, 1 μM Yoda1: 32.3 ± 11.0, 5 μM Yoda1: ± 12.1)(10^−18^ N × m). The data from the inset are shown in Figure 7E.

**Video 1**

RBCs in complete medium were placed in poly-L lysine coated slide chamber (Ibidi μ-slide VI). The chamber was then flushed with complete medium containing 5 μM Yoda1 and time–lapse videomicroscopy recorded at 37°C with 63 x magnifications at a speed of 1 image/4 seconds. Time in sec indicated relative to the addition of Yoda1. Scale bar 10 μM.

**Video 2**

To observe the recovery of cell shape, RBCs in complete medium were treated with 10 μM Yoda1 (at this concentration, all RBCs become strong echinocytes), directly placed in glass bottom dishes (MatTek), and time–lapse videomicroscopy recorded at 37°C and 5% CO2 with 63 x magnifications over 2 ½ h. Time indicated in min. Scale bar 10 μM.

**Video 3**

Representative video of invasion events with mock-treated RBCs. RBCs that become invaded are labelled 1 and 2. Selected images of RBC 1 is shown in Figure 7.

**Video 4**

Representative video of the reaction of RBCs pretreated with 5 μM Yoda1 to contact with *P. falciparum* merozoites and absence of invasion. Scale bar 10 μM.

